# Alveolar early progenitors in the aged human lung have increased expression of ACE2 accompanied with genes involved in beta-amyloid clearance: Indication of SARS-CoV-2 also using soluble ACE2 in aged-lungs to enter ACE2-negative cells

**DOI:** 10.1101/2020.05.25.115774

**Authors:** Virendra K. Chaudhri

**Affiliations:** Department of Immunology, University of Pittsburgh, 200 Lothrop Street, Pittsburgh, PA, USA, 15213

## Abstract

COVID-19 is the current pandemic caused by severe acute respiratory syndrome virus 2 (SARS-CoV-2) that uses ACE2 protein on the cell surface. By analyzing publicly available datasets, I uncovered that alveolar early progenitors (AEP), a subset of the type-2 pneumocytes, showed increased ACE2 expression in the older lungs. AEPs co-express TMPRSS2, CTSL. Aged AEP-gene expression signature suggested an active response to beta-amyloid-induced ACE2 shedding, to limit the intercellular beta-amyloid accumulation in otherwise healthy human lungs. Susceptibility of AEP to SARS-CoV2 and ACE2 secretory capacity of these cells makes aged human lung sensitive for rapid-infection, by a possible in-solution ACE2 binding and entry into ACE2-negative cells, thereby increasing the target cell diversity and numbers. Single-cell analysis of COVID19 patients with moderate and severe infections, clearly showed that severe infections showed SARS-CoV-2 transcript in ACE2-negative TMPRSS-negative but CTSL-positive cell types in their bronchoalveolar lavage fluid, validating in-solution ACE2-binding enabling infection.

---

Since December 2019, SARS-CoV-2(COVID-19) infection, which was first reported in Wuhan China (Zhu et al., 2020), has rapidly spread and turned into a worldwide pandemic, claiming as of May 25, 2020, more than 5.4 million infections with more than >344,000 deaths. COVID-19 has ~3-4% fatality rate (https://coronavirus.jhu.edu/map.html). COVID-19 virus belongs to the family of MER viruses, which causes infections in the human respiratory system leading to severe respiratory syndrome that can result in death. More importantly, the SARS-CoV-2 develops atypical pneumonia with a higher risk for older individuals or individuals with lung, heart, and kidney-related health issues (Shen et al., 2020), leading to a higher fatality rate accompanied by rapid progression of the respiratory syndrome and atypical pneumonia (Huang et al., 2020). SARS-CoV-2 binds to angiotensin-converting enzyme-2 (ACE2) binding followed by priming of spike protein S2 by transmembrane serine protease-2 (TMPRSS2) to enter the cells. Once internalized, Cathepsin L activates spike protein S, allowing the fusion of endosomal and viral membranes. A recent report has identified several epithelial cell subsets type-II pneumocyte, club cells, and ciliated cells in the lungs have ACE2, TMPRSS2 and CTSL expression. These are key genes for establishing COVID-19n infection(Hoffmann et al., 2020).

Therapeutic interventions of SARS-CoV-2 needed a biological basis and possibly a mechanism of its severity. I used an analytical approach to integrate public datasets to answer the question, why the aged human population is at risk of severity and a higher fatality rate by SARS-CoV-2 ? The analysis is centered around the essential genes ACE2, TMPRSS2 and CTSL, that encodes the proteins implicated and shown to be responsible for entry to establishing the viral infection. The analysis focuses on isolating any subset of cells that are potentially altered in aged-human lungs to support SARS-CoV-2 and what is the nature of this alteration that can give insights into therapeutic intervention.

To isolate cells particularly susceptible to SARS-CoV2, I used public datasets (see methods) from the scRNA-seq profiles of lung alveoli of 6 individual donors, where three donors were less than 30 years in age, and three were 55 years or older (Reyfman et al., 2019). Using the Seurat suite in R, the data was processed to remove batch effects. Cell types were separated using the Louvain clustering algorithm (see Methods for details) (Supplementary Data Table S1). Clustering identified 13 cell types in single-cell analysis and are shown in Figure 1 A as UMAP projections (Supplementary Table S1A). While ACE2 expression was primarily restricted to AT2 and Ciliated cells, TMPRSS2 was expressed in AT2, AT1, Club cells, Ciliated cells, and several immune subsets (Fig1 B-C). ACE2 and TMPRSS are required by SARS-CoV2 to enter the cell, while CTSL is needed to establish infection once the virus has entered the cell (Fig1.B-C)(Bosch et al., 2008). While ACE2 and TMPRSS are more expressed among pneumocytes, the CTSL expressed at moderate levels in pneumocyte and higher expression in macrophages, monocytes and other immune cells (Fig1. B-C). Given these data, the susceptibility map of lung cell populations for SARS-CoV-2 was limited for a very small subset in AT2 cells in lung alveoli that expressed ACE2, TMPRSS2 and CTSL.

**Figure 1:**
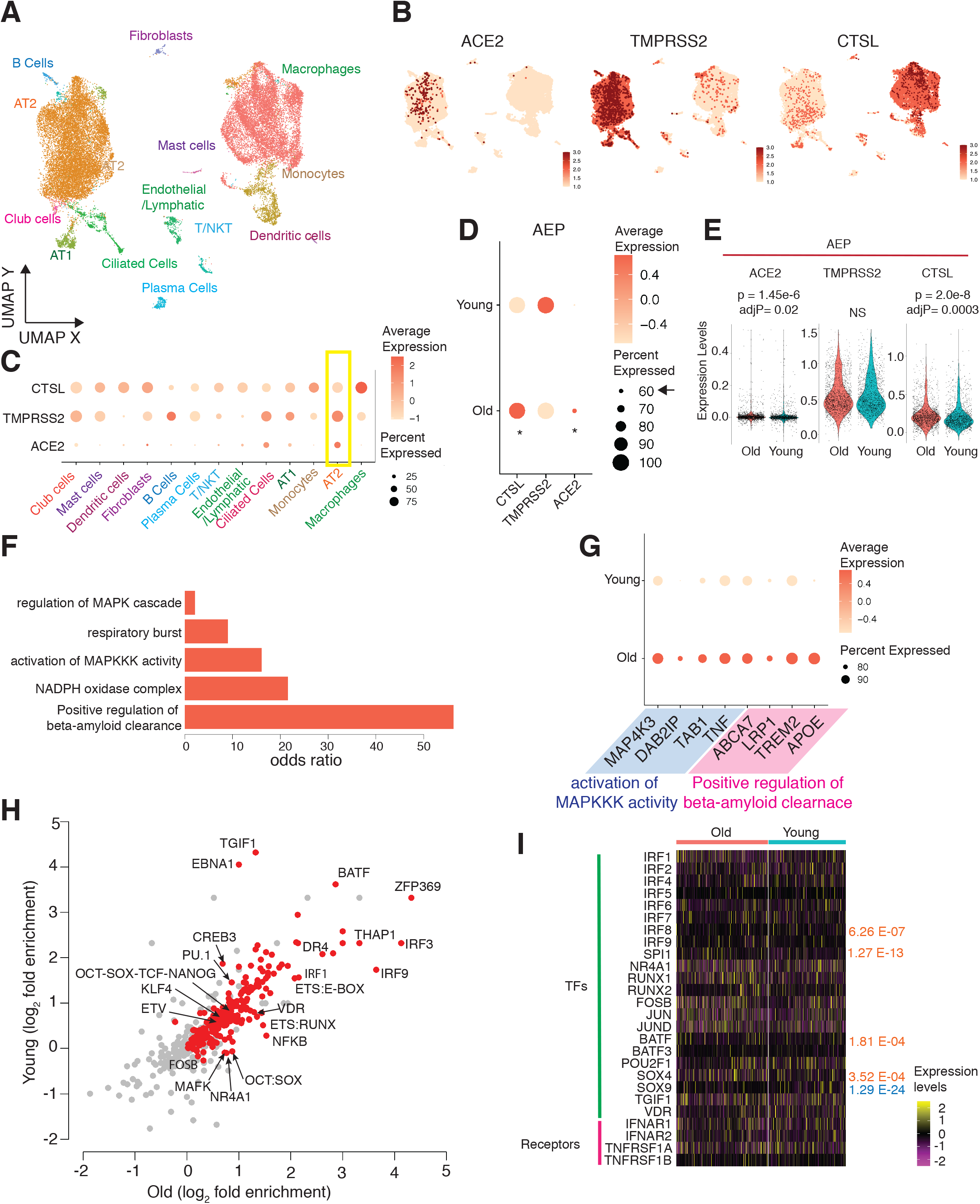
scRNA seq analysis of human lung alveoli reveals alveolar early progenitors (AEP) with age-associated upregulation of ACE2 correlated with beta-amyloid clearance genes to suggest active renin-angiotensin signaling (RAS): **A**. The scRNA-seq libraries from ~30,000 cells as Louvain clusters and identified cell types are plotted on uniform manifold approximation (UMAP) co-ordinates. Cells are color-coded for cell types. **B**. UMAP projections with expression levels of critical genes for SARS-CoV-2 ACE2, TMPRSS2, and CCTSL are shown as cell level heatmaps. **C**) a dot plot showing the viral entry genes expression across all identified populations. A yellow box indicates AT2 cells with both higher levels and a high percentage of cells expressing these genes. **D-E**. Dot plot and violin plots, respectively, showing the expression levels layered for the genes in B in AEP cell types. **F.** The odds ratio for key GO terms from GO enrichment analysis for the marker genes for old-AEPs is plotted here. **G.** genes from 2 key GO terms are plotted as dot plots comparing old vs. young AEPs. **H.** TF motif enrichment analysis of promoters of old-AEP vs. young-AEP genes is plotted here as log2 fold enrichment over scrambled background sequences for two groups as scatter plot. Red dots indicate enrichment with logP value less than −10 in either group. **I.** Heatmap showing relative expression levels of selected transcription factors and cell surface receptors in old-AEPs and young-AEPs.

I further focused on AT2 cells to isolate a subset recently identified as alveolar early progenitors (AEP) among AT2 cells(Zacharias et al., 2018). Bulk RNA-seq data were obtained from (Zacharias et al., 2018) and used to generate the expression profile of the human AEPs by averaging the replicate log2 FPKM values. AT2 cells with correlation value 0.5 or higher to AEP expression profile were assigned as AEPs(n=2948). Cells other than AEPs were identified as AT2(11,076) cells and therefore referred as such in the text now onwards. The AEP subset was further segregated based on the age of donors as young donors (n=3, age < 30years), termed young-AEP, and old donors (n=3, age >55 years, old AEP) termed old-AEP. When compared the expression profiles of old and young AEPs, old-AEPs showed a remarkable increase in both ACE2 expression and percentage of cells expressing AEP2 (Fig1.D-E). A similar analysis of AT2, AT1 and Ciliated cells did not show such a relationship (Supplementary Figure S1, A-C). TMPRSS2 shows no significant change, and CTSL levels were increased in older AEPs (Fig 1D-E).

Current results are consistent with the observation of SARS-CoV2 prevalence in older individuals and the first indication of why and which cell subset may be primarily contributing to the SARS-CoV2 pathogenesis of aged lungs. Since AEPs are the cells that respond rapidly to lung injury and rapidly differentiate into AT2 cells and AT2 then give rise to AT1 and Ciliated cells, a slingshot analysis on scRNA-seq revealed that this trajectory was successfully isolated from scRNA-seq data set (Supplementary Figure S2, D). Independently Bulk RNA-seq of AEP and AEP depleted AT2 cells showed that ACE2 is detected only in AEP when compared to AEP depleted AT2 cells (p value=0.035, determined by cuffdiff (Trapnell et al., 2010), also see Supplementary Data Table 2). This further validated AEP identity and concentration of ACE2 in this subset.

Next, to isolate how these AEPs are changed in the aged lung, marker genes were identified for old and young AEPs (see methods for details). Marker genes for old-AEP and young-AEP were identified with criteria being in the top quartile of the percentage change in the number of cells expressing them and in the top quartile of log fold change with an adjusted p-value of less than 0.05. This revealed 532 genes as old-AEP markers and 220 genes as markers of young-AEPs(Supplementary Table S1B-D).

A hypergeometric gene ontology (GO) testing revealed a distinct expression signature for the aged AEPs and is summarized in (Figure 1. F and Supplementary Table 1A-B). Surprisingly genes related to beta-amyloid clearance and MAPK pathway genes, including TNFα and MAP4K3, were upregulated in old-AEPs. Beta-amyloid clearance is a well-studied process in relation to Alzheimer’s in the human brain, where several are proteases and peptidase are secreted, in response to extracellular beta-amyloid accumulation (Panza et al., 2019; Yoon and Jo, 2012). ACE2 deficiency is also known to increase the risk of early onset of Alzheimer’s. (Kehoe et al., 2016).

More importantly, some of the genes which were upregulated in old AEPs are also known for their association with Alzheimer’s and considered marker genes for Alzheimer’s risk, namely APOE, TREM2, LRP1 and ABCA7 (Fig1. G)(Karch and Goate, 2015). APOE is a lipoprotein that binds extracellular amyloids to be internalized in TREM2 dependent manner, a pathway that is sensitive to hydroxychloroquine (Jensen et al., 1994; Yeh et al., 2016), an antimalarial medication recently tested for COVID-19 intervention (Geleris et al., 2020). While lung amyloidosis has several subtypes, it is plausible that these genes are upregulated in response to amyloid accumulation in aged lungs as risk factor increases for lung amyloidosis with age (Milani et al., 2017) and amyloid accumulate linearly with age (Gonneaud et al., 2017).

Further, genes for the NADPH complex were enriched in old-AEPs (Figure 1F), and it is known that renin-angiotensin signaling involves increase NADPH activity accompanied by increased mitochondrial activity(Xu et al., 2011). It was no surprise to see genes for respiratory burst, which indicates the rapid generation of reactive oxygen species (ROS) (Figure 1F) as an active RAS system involves rapid ROS generation. As angiotensin signaling requires either TypeI or Type2 angiotensin receptor transcripts for genes of either receptor can validate an active RAS system. Type2 angiotensin (AGTR) receptor was detected in both young and old AEPs, while levels were significantly higher in old-AEPs (Figure S1E, p=1.02e-12). No transcript for type1 receptors genes AGTR1 was detected in AEPs. Since AGTR2 is known to present earlier in development and expression in adult tissue is rare, an increase in the expression of AGTR2 transcripts in old-AEP further attested the active RAS.

The renin-angiotensin signaling system has been studied in the heart, lung, brain and other tissue types; it has not addressed the function of ACE2 and CTSL in AEPs. Angiotensin II receptor signaling is known to causes ACE2 shedding in human airway epithelia (Jia et al., 2009). Further, the current analysis also showed that in older lungs, AEP and Macrophages both have TNFα levels significantly increased (Figure 1G, Supplementary Figure S1F). Along with the gene expression signature described above, it is plausible aged baseline inflammation very likely be a result of the increased beta-amyloid load (Decourt et al., 2017). ADAM17 member of "A Disintegrin And Metalloprotease" (ADAM) family, also known as TNFα converter (TACE), mediates ACE2 shedding on the membrane in response to RAS mediated signaling (Lambert et al., 2005), was expressed in AEPs (Supplementary Figure S1G). ADAM17 expression levels did not show any age-related changes in AEP (Supplementary Table 1B). Together this data argues that changes in AEP in older individuals are reflective of response to angiotensin signaling that results in significant transcriptional changes observed here. This also suggest in aged lungs then very likely have a higher concentration of soluble and catalytically active ACE2.

Given that now it was established that in aged lung AEPs’ expression profile suggestive of activation of RAS system in response to amyloid accumulation and likely shedding ACE2, I further extended the analysis to promoter regions (−1000, TSS, +100bp) of genes that were either upregulated (old genes) or down-regulated (young gene) in AEPs. Promoter regions were scanned for old and young AEP genes separately with a corresponding 10-fold set of scrambled sequences of test sequences with preserved dinucleotide frequencies as described previously (Chaudhri et al., 2020). Both sets of promoters enriched for stem cell TF motifs, namely OCT, SOX, NANOG, and AP1 family members. Promoters of genes upregulated in old-AEPs enriched for motifs for IRF, NFKB, ETS, VDR(Vitamin D Receptor), while promoters of genes that were downregulated in aged-AEP were enriched for motifs for TGIF, CREB3 and PU.1. It has been recently shown that ACE2 levels in human lung pneumocytes upregulate in interferon dependent manner (Ziegler et al., 2020); notably, in Alzheimer’s amyloid induced neuro-inflammation is driven by type-1 interferons(Roy et al., 2020). Another strong parallel with microglia like function by AEPs and AEPs expressed interferon receptors as well as receptors for TNFα as well as these transcription factors (Figure1.I). This analysis reveals that the upregulation of these genes is very likely a result of amyloid accumulation mediated RAS-angiotensin signaling and localized TNFα and interferon-mediated inflammation. This analysis further validated that these genes are a coherent set of genes that can have similar responses to further increase to interferon, and ACE2 expression increase will be much higher in antiviral response. Additionally, active RAS is very likely contributing to ACE2 shedding via ADAM17, thereby having a high local concentration of soluble ACE2 in intercellular space. Based on this analysis, a plausible hypothesis is unlike SARS-CoV, SARS-CoV-2 can potentially bound to soluble ACE2 and can then bind to TMPRSS2 positive cells to prime spike protein S and can enter independent of membrane ACE2 expression. This explains the severity of SARS-CoV-2 infections and its correlation with age.

To validate the hypothesis above from the analysis of healthy lung alveoli scRNA-seq data and show that soluble ACE2 can be a reason for the severity of SARS-CoV-2 infection older people, I used a recently published Broncho-alveolar lung fluid (BALF) scRNA-seq from human subjects with confirmed SARS-CoV-2 infection (Liao et al., 2020). This data is unique as transcripts were also mapped to the SARS-CoV-2 genome and termed "nCoV". 10X data files were processed and analyzed similar to the analysis shown above, and Louvain clustering of cells in BALF is shown in Figure 2A. No AT2 or AEP cells were detected in BALF samples, and only identified epithelial cell types present were ciliated and club cells. Surprisingly when expression levels of ACE2, TMPRSS2, CTSL and nCoV (RNA reads mapped to COVID-19 genome) overlaid on UMAP projections, it showed nCoV expression in cells which do not express ACE2 or TMPRSS2, yet these cells had high levels of CTSL (Figure 2B). This strongly suggested soluble ACE2 once bound to SARS-CoV2, the necessity of surface TMPRSS2 becomes redundant for priming spike protein and either some other transmembrane serine proteases can prime the spike protein on cell surface receptor or in solution secreted proteases.

**Figure 2:**
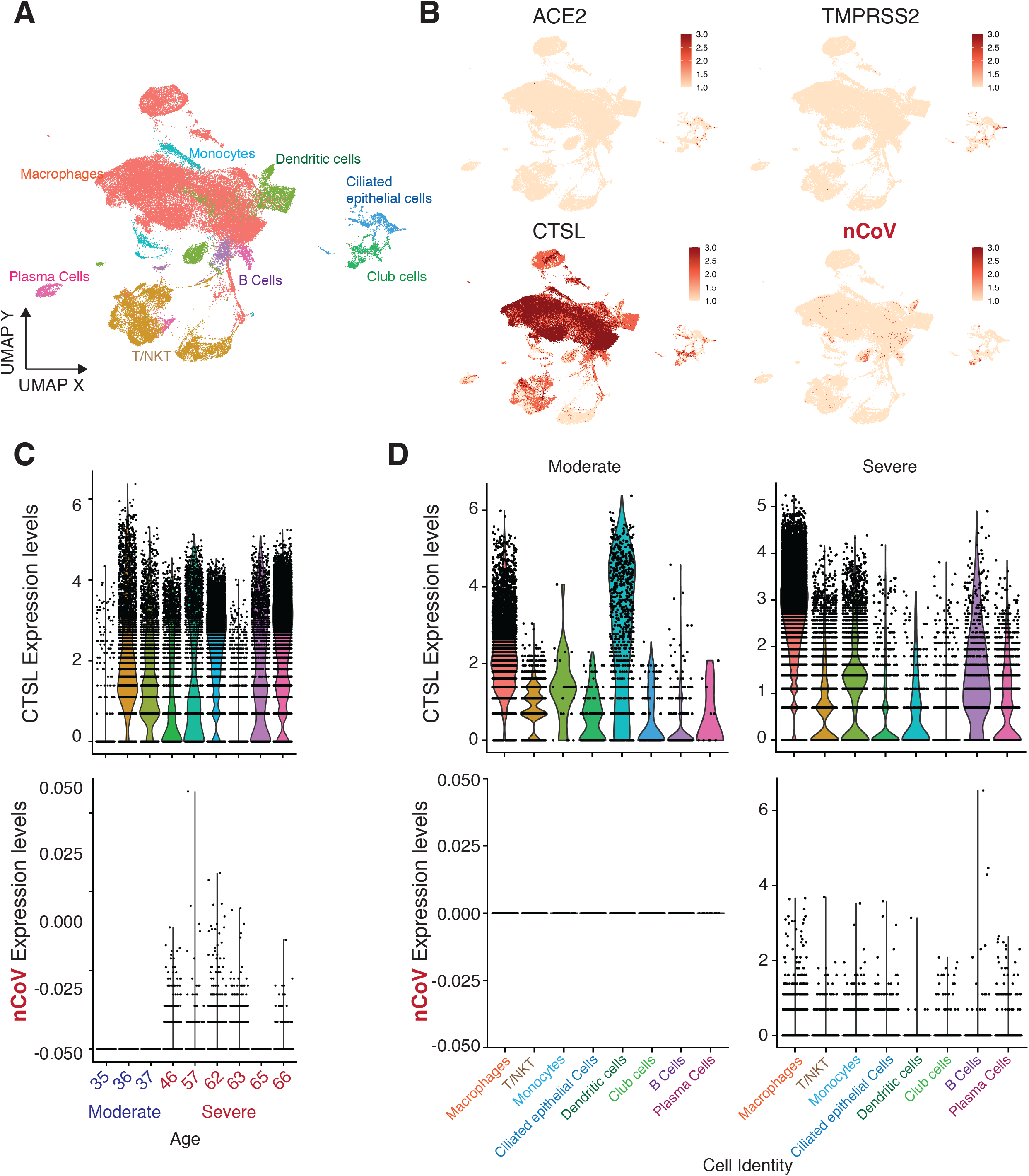
Validation of SARS-CoV-2 infection in ACE2-neg and TMPRSS2-neg cells. **A.** Louvain clustering of ~53,000 cells in bronchoalveolar lavage fluid (BALF) from moderate and severely infected subjects showing identified cell types as indicated, on UMAP co-ordinates. **B.** UMAP feature plot of genes responsible for viral infection alongside nCoV transcripts, which are the RNA-seq reads mapping to the SARS-CoV-2 genome. **C**. Violin plots showing CTSL and nCoV transcripts across different cell types in BALF from moderate and severely infected groups.

In-solution binding to ACE2 and TMPRSS2 independent priming explains why CTSL high cells showed nCoV transcripts seemingly across many cell types. When compared to the CTSL and nCoV transcript levels and relationship with age of the subjects, severe cases were older individuals, and nCoV transcripts were only detected in older human subjects, while CTSL levels varied across both moderate and severe groups and did not show any age association (Figure2C). ACE2 and TMPRSS2 profile ACE2 only detected in older(severe) subjects while TMPRSS2 was detected in all subjects. The next question comes, which of the cell types in BALF are infected, and the presence of nCoV transcripts can confirm that. As hypothesized from the analysis of healthy human lung alveoli, in-solution SARS-CoV-2 binding, expectedly all cell types in the severe group expressed varying levels of nCoV transcripts (Figure 2D). ACE2 and TMPRSS in the severe group were primarily expressed in epithelial cells and TMPRSS2 and not other cell types. The presence of nCoV transcript across several cell types, including lymphocytes, explains the complications of SARS-CoV-2 infection in older people. This result is the opposite of the current dogma of SARS-Co-2 entry solely through membrane receptors. These data strongly argued that SARS-CoV-2 has altered surface structure with the ability to use soluble ACE2, and then infect cells independent of surface ACE2 or TMPRSS2 expression.

In line with the analysis of healthy lung alveoli, that ACE shedding by AEPs in the aged-lungs are providing a higher localized concentration of ACE2 in alveoli. It is reasonable as entry of SARS-CoV2 into cells by priming of S protein once bound to ACE2 can occur independently by TMPRSS2. Alternatively, this priming can be occurring on the membrane of TMPRSS2 expressing cells but not necessarily entering the same cell thereby TMPRSS2 positive cells acting as both virus recipient and enabling priming of the virus to infect cells independent of ACE2 or TMPRSS2 receptor expression, strongly supported by Figure 2D. Another possibility is some other secreted serine proteases can do the priming once soluble ACE2 is bound to SARS-CoV2. It is important to note that COVID-19 has substantial changes in its surface that support virus survival differently than its SARS-CoV-1 from 2003(van Doremalen et al., 2020), and that might attribute to changes in binding preference to ACE2.

It has been reported that COVID-19 infection is severe for people with higher age, diabetes, heart conditions, and in many of these conditions increase in ACE2 shedding has been studied, which adds to the validation of current proposed model. It is important to note that even small amounts of secreted ACE2 in the alveoli niche can have very high local effective concentrations for binding by SARS-CoV-2. Recently it has been shown that there is a 3 fold lower mortality rate from SARS-CoV-2 in individuals on medication with ACE2 or AT2-receptor (AGTR2) inhibitors for hypertension, compared to those who were not on those medications in that group (Zhang et al.). ACE2 inhibitors used in hypertension or other cardiovascular issues were recently considered therapeutic interventions but have been criticized for no strong medical basis. With the current study proposed, it will be beneficial to use these inhibitors but directed to the lung, and it can lower the infection rate by SARS-CoV-2. Recently a clinical trial of using soluble recombinant hACE2 was attempted, which was withdrawn (NCT04287686). Finally, the analysis presented here strongly supports that altered surface structural changes of SARS-CoV-2 might contribute to the altered preference for ACE2 binding, and different modes of transmission. These findings suggest a therapeutic route that interferers with soluble ACE2 binding or inhibition of ACE2 secretion in lungs may be the fastest acting path.

Future work towards experimentally detecting in-solution ACE2 bound SARS-CoV-2 and its ability to infect cells not expressing ACE2 will further the advances in designing suitable therapeutics.

## Methods

### scRNA-seq data analysis

Raw 10X counts data for human lung alveoli was obtained from (GSE122960) as individual 10X output files from a published study (Reyfman et al., 2019). Data from active smokers was discarded. All data processing was performed by importing in R using Seurat (Stuart et al., 2019), and several Seurat routines were used to downstream analysis. Cells were selected based on the minimum expression of 200 genes, and genes are selected to be expressed in at least 5 cells in each of the individual samples. Cells with more than 10% mitochondrial gene expressed were excluded. For all 6 non-smokers, the count data was processed individually using Seurat function SCTransform with 12,000 variable features. In final selection Cells were selected with at least 500 expressed genes and less than 7500 expressed genes to avoid doublets. This data was then used to identify up to 15,000 anchors using function “FindIntegrationAnchors” across samples. Finally, data from all six samples were integrated using function “IntegrateData,” and all gene features were integrated. Dimension reduction was done using PCA in Seurat using 100 components. Cell populations were clustered using Lauvain clustering implementation in Seurat by using all 100 PCs. marker genes identified for each of the cluster from using Seurat function “FindAllMarkers”(Supplementary Data Table S1). To identify the cells, marker genes were obtained from (website and reference) and intersected with identified marker genes. These markers were used to comprehensibly identify the populations using the marker genes from previously published source study and additionally verified from the markers using CellMarker database (Zhang et al., 2019). Marker genes for old AEP and young AEP were identified with meeting 3 criteria 1) criteria being in top quartile of percentage-change in cells expressing them 2) in top quartile of log fold change with 3) an adjusted p-value of less than 0.05 determined by function FindMarkers. All plots are generated in R using R-base function or Seurat’s visualization functions. For slingshot Seurat UMAP co-ordinates were used with default settings and AEP as a starting point (Street et al., 2018).

BALF scRNA-seq obtained from (GSE145926) and analyzed as stated above with the exception of while selecting the cells for SCTransform minimum expressed genes were set to 200 and maximum genes at 6,000 in line with the source study (Liao et al., 2020).

### Bulk RNA-seq expression signatures of human AEP

Bulk RNA-seq data for human AEP is obtained from an earlier study (Zacharias et al., 2018)(SRA id for AEP samples: SRR5379121, SRR5379122, SRR5379123; SRA ids for AEP depleted AT2: SRR5379118, SRR5379119, SRR5379120). Fastq files were aligned to hg19 genome assembly using STAR(Dobin et al., 2013) and transcript abundances were determined by Cufflinks default options (Trapnell et al., 2010). Of these based on PCA, sample correlation and very high levels of AEP marker "TM4SF1" in sample SRR5379120, it was excluded from AEP depleted AT2 group for further analysis. Resulting data were log-transformed and the mean expression value of each gene is used to calculate the Pearson correlation coefficient of AT2 cells in scRNA-seq. This correlation was calculated by using only genes present in scRNA-seq and bulk RNA-seq. The correlation coefficient value of any AT2 cell with AEP expression profile with a value 0.5 or higher is assigned as AEP. cuffdiff function from cufflinks was used to generate differential gene expression between AEP and AEP depleted AT2 cells.

### Gene Ontology enrichment of genes and TF motif enrichment analysis

Gene ontology (GO) enrichment and promoter TF motif analysis is performed as described earlier(Chaudhri et al., 2020). GOstats package (Falcon and Gentleman, 2007) was used for hypergeometric testing and 15,080 genes in integrated scRNA-seq data were used as gene universe. For TF motifs analysis, protein-coding marker genes annotated in hg19 genome assembly were used. Promoter sequences of marker genes (−1000bp, TSS, +100 bp) were used as test set, and a 10-fold scramble sequence set of these test sequences is generated for the background set. Homer functions “homer2 known” with default settings were used for motif enrichment analysis (Heinz et al., 2010).

---

**Supplementary Figure S1:**
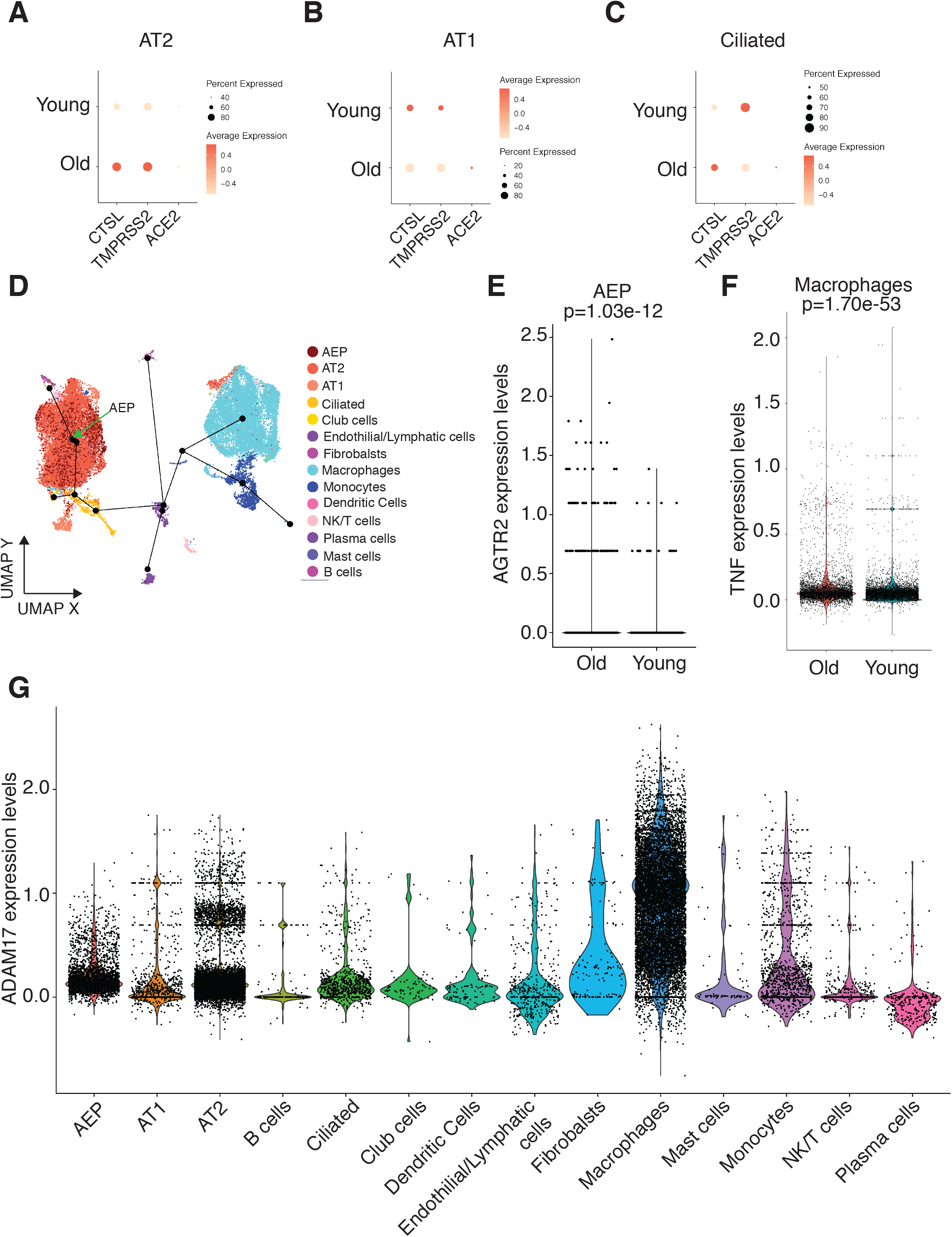
Age-associated upregulation of ACE2 is limited to AEP and suggest AEPs as the sole source of shedded ACE2, related to Figure 1: **A-C** shows dot plots of ACE2, TMPRSS2, and CTSL levels in AT2 (A), AT1(B) and Ciliated(C) cells for age-associated changes in the expression and number of cells expressing them. None of the changes were significant three cell types. **D.** Slingshot trajectory analysis is plotted; the black line represents trajectory starting at AEP (indicated by the green arrow), with the following dots representing cell types. **E.** old-AEPs express higher levels of transcripts for AGTR2, a type 2 angiotensin receptor. Violin pot for expression levels for the AGTR2 gene in old and young AEPs is plotted here. P-value is from the Wilcoxon rank-sum test. **F.** TNFα levels in macrophages in old and young AEPs are shown as violin plots. Wilcoxon rank-sum test is used to calculate the p-value. **G.** The expression of ADA17 (TNFα maker) is plotted across all cell types as a violin plot, including AEP. AEP, AT2, and macrophages primarily expressed ADAM17.

**Supplementary Figure S2:**
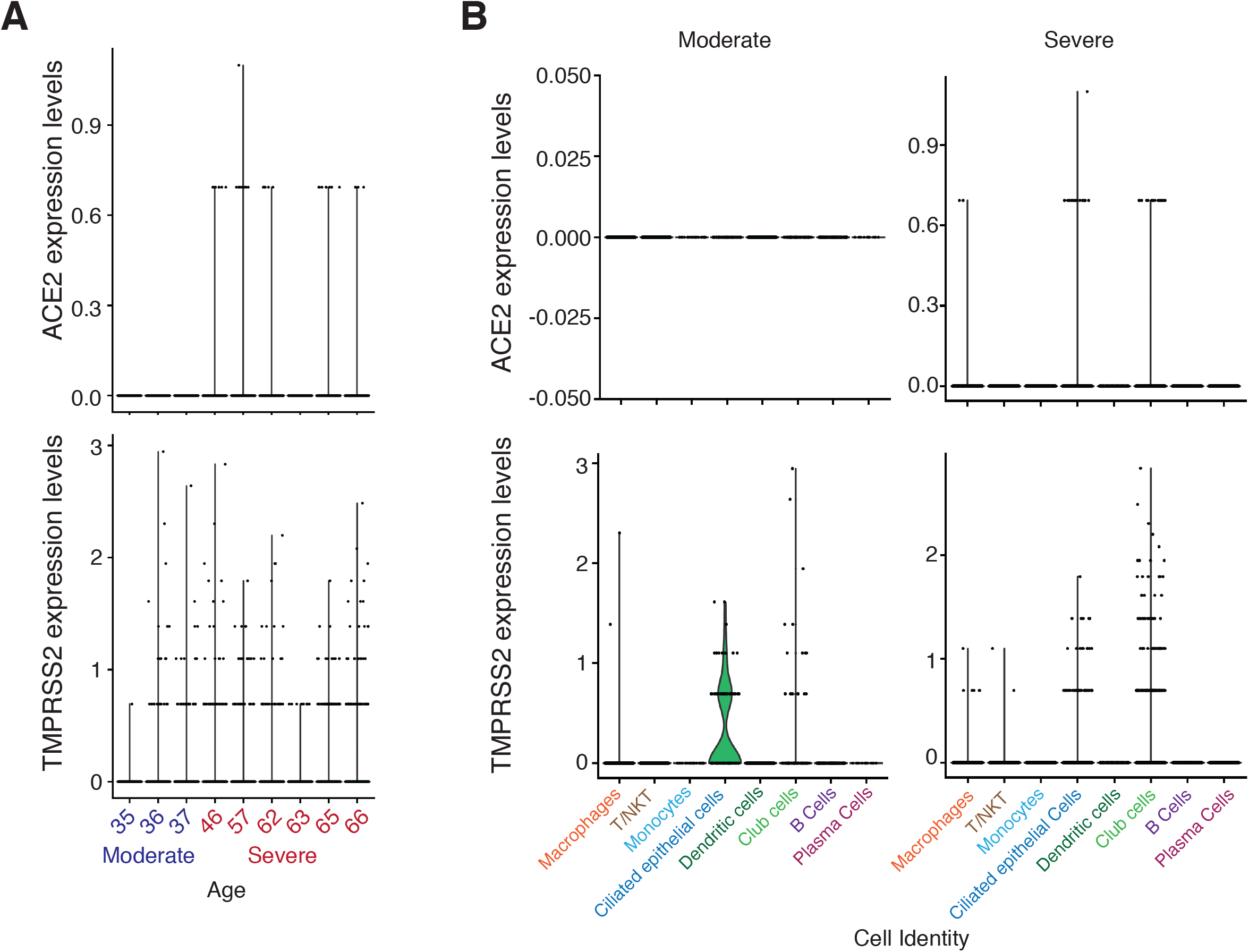
Lack of ACE2 and TMPRSS2 does not limit the SARS-CoV-2 infections, related to Figure 2. **A.** ACE2 and TMPRSS2 expression levels of individual patients with SARS-CoV-2 are plotted as violin plots, where the x-axis indicates the patients’ age**. B.** Violin plots showing ACE2, and TMPRSS2 levels in different cell types present in BALF scRNA-seq.

